# Venlafaxine stimulates PNN proteolysis and MMP-9 dependent enhancement of gamma power; relevance to antidepressant efficacy

**DOI:** 10.1101/432419

**Authors:** Alaiyed S., Bozzelli P. L., Caccavano A., Wu J.Y., Conant K.

## Abstract

Drugs that target monoaminergic transmission represent a first-line treatment for major depression. Though a full understanding of the mechanisms that underlie antidepressant efficacy is lacking, evidence supports a role for enhanced excitatory transmission. This can occur through two non-mutually exclusive mechanisms. The first involves increased function of excitatory neurons through relatively direct mechanisms such as enhanced dendritic arborization. Another mechanism involves reduced inhibitory function, which occurs with the rapid antidepressant ketamine. Consistent with this, GABAergic interneuron-mediated cortical inhibition is linked to reduced gamma oscillatory power, a rhythm also diminished in depression. Remission of depressive symptoms correlates with restoration of gamma power.

Due to strong excitatory input, reliable GABA release and fast firing, PV neurons represent critical pacemakers for synchronous oscillations. PV neurons also represent the predominant GABAergic population enveloped by perineuronal nets (PNNs), lattice-like structures that localize glutamatergic input. Disruption of PNNs enhances lateral diffusion of glutamate receptors, reduces PV excitability, and enhances gamma activity.

Studies suggest that monoamine reuptake inhibitors reduce integrity of the PNN. Mechanisms by which these inhibitors reduce PNN integrity, however, remain largely unexplored. A better understanding of these issues might encourage development of therapeutics that best upregulate PNN modulating proteases.

We observe that the serotonin/norepinephrine reuptake inhibitor venlafaxine reduces PNN integrity in murine brain. Moreover, venlafaxine treated mice (30 mg/kg/day) show an increase in carbachol-induced gamma power in hippocampal slices. Studies with mice deficient in matrix metalloproteinase-9 (MMP-9), a protease linked to PNN disruption in other settings, suggest that MMP-9 contributes to venlafaxine-enhanced gamma activity.

## Introduction

Major depressive disorder (MDD) is a debilitating condition that affects ∼12–17% of individuals in the U.S.at some point during their lifetime. This places a substantial burden on those affected and on family and friends. Moreover, untreated depression may increase one’s risk for substance abuse and Alzheimer’s disease (1, 2).

While a full understanding of molecular underpinnings is lacking, studies suggest that depression is linked to reductions in the strength of glutamatergic synaptic subsets in regions including hippocampus and prefrontal cortex (PFC). Animal models have shown changes including reduced GluA1 expression in layer V pyramidal cells, and impaired hippocampal long-term potentiation (LTP) [reviewed in (3)].

In terms of effective treatment, one mechanism by which antidepressants may enhance glutamatergic transmission involves generation of new glutamatergic synapses. Animal models support this as successful treatment has been linked to an increase in the number of dendritic spines, which represent post-synaptic processes for the majority of excitatory synapses in the central nervous system (4). Enhanced dendritic spine number could potentially follow from antidepressant-mediated increases in neurotrophins.

A non-mutually exclusive mechanism by which primary or adjunct therapeutics could improve glutamatergic signaling is through reduced inhibition of glutamatergic neurons. This possibility is supported by the rapid antidepressant activity of ketamine, which can stimulate disinhibition by acting as a preferential antagonist for GluNs localized to inhibitory interneurons (3). Importantly, new research has shown that negative allosteric modulation of the α5 GABA-A receptor subunit increases electroencephalographic gamma power (5), a rhythm that is reduced in human depression and an animal model of depression, and also normalized with remission (6, 7). The same allosteric modulator concomitantly improves performance in the forced swim test (5).

An alternative means to affect cortical disinhibition is through manipulation of the perineuronal net (PNN), a lattice like extracellular matrix that is predominantly localized to parvalbumin (PV) expressing interneurons (8, 9). The PNN facilitates properly localized glutamatergic input to PV cells (10), thus enhancing PV mediated inhibition. Disruption of the PNN leads to increased lateral diffusion of glutamate receptor subunits on PV cells (10), and it may also result in increased diffusion of glutamate. Consistent with this, enzymatic degradation of the PNN reduces the frequency of excitatory post synaptic currents in hippocampal fast spiking interneurons (11). Recent work also suggests that PNN disruption can enhance cortical excitation and gamma activity (12).

In terms of antidepressant therapy and the PNN, it is of interest that chronic treatment with fluoxetine decreases PNN staining in murine medial PFC and hippocampus (13). Recently published work has also shown that a stress related model of depression increases PNN staining (14).

A relatively unexplored question is whether PNN reduction makes a substantial contribution to antidepressant efficacy and, if so, whether therapeutics or combinations beyond fluoxetine may best achieve this. The significance of this possibility is underlined by the widespread distribution of PNN enwrapped PV interneurons. As opposed to GABA-A α5 subunits, which have a depression-relevant but relatively restricted expression in forebrain, PNN enveloped PV neurons are found in multiple brain areas important to mood and memory.

Studies that have examined PNN disruption for its potential to mediate cortical disinhibition have typically employed hyaluronidase, which targets the PNN backbone, or chondroitinase, which targets chrondroitan sulfate proteoglycan side chains. Physiologically relevant modulators of PNN integrity are, however, expressed in the central nervous system. Important among these modulators are proteases belonging to the metalloproteinase (MMP) family.

MMP-9, MMP-13, and select transmembrane MMPs have been well linked to PNN component cleavage *in vivo* (15). For example, light reintroduction after dark exposure triggers MMP-9 dependent PNN degradation (16). In addition, seizure activity has been shown to increase MMP-9 levels and PNN breakdown while pretreatment with a broad-spectrum MMP inhibitor abrogates these effects (17). Further, in murine models of brain injury, selective MMP inhibition also reduces PNN remodeling (18).

Of interest, PNN degrading MMPs may be upregulated by monoamines or reuptake inhibitors (19). Consistent with this possibility is work demonstrating increased MMP-9 expression in venlafaxine treated rats (20). While effects of specific monoamines on MMP release from brain-derived cells have not been extensively studied, in non-neural cells, norepinephrine has been shown to enhance MMP expression as well as MMP dependent endpoints (21).

In the present study, we focus on the question of whether one of the most commonly prescribed antidepressants, venlafaxine, can stimulate PNN proteolysis and disinhibition as measured by its ability to upregulate carbachol-induced gamma power. Our studies focus on the hippocampus, a region increasingly implicated in depression (22). We also explore a role for MMP-9 in venlafaxine-stimulated disinhibition.

## Materials and Methods

### Chemicals and reagents

Venlafaxine and carbamoylcholine chloride (carbachol) were purchased from Sigma Chemical (St. Louis, MO; catalogue numbers PHR1736 and C4382). Carbachol was maintained in frozen 50 mM aliquots and reconstituted in ACSF to a final concentration of 40 μM just prior to slice treatment.

### Subjects and drug administration

Both male and female C57BL/6J (Jackson Laboratory, #000664) or MMP-9 homozygous null mice (Jackson Laboratory, #007084) that have been backcrossed to C57BL/6J mice for at least five generations were used for experiments. Both mouse strains were bred in-house. At approximately 4 weeks of age, a time at which hippocampal PNN deposition is complete (18), mice were treated for 2 weeks with saline or saline containing 30 mg/kg venlafaxine. While lower doses act predominantly on serotonin reuptake, this is a dose that has been shown to reduce uptake of both serotonin and norepinephrine (23). Treatments were delivered in a total volume of 200 μl and administered daily by intraperitoneal injection. Experiments were performed in accordance with National Institutes of Health guidelines and an institutionally approved protocol.

### Preparation of brain lysates and ELISA

Following saline or venlafaxine treatment and euthanasia, hemi brains were dissected and hippocampi were extracted and lysed in RIPA buffer [50 □ mM Tris, pH 7.5, 150 □ mM NaCl, 0.1% SDS, 1% IGEPAL, and 1X protease and phosphatase cocktail (Thermo Scientific 1861281)]. Lysates were sonicated for 10 □ seconds, placed on ice for 20 □ minutes, and centrifuged for 15 □ minutes at 14,000 □ rpm in 4 □ °C. Lysate supernatants were saved for analysis. Pro MMP9 protein concentration in hippocampal lysates were measured by ELISA, performed according to the manufacturer’s protocol (Mouse Pro-MMP-9, R&D systems P162135).

### PNN staining and microscopy

Following saline or venlafaxine treatment and euthanasia, hemi brains were fixed overnight in 4% PFA/sucrose at 4° C. Brains were then paraffin-embedded and sectioned at 15 microns. Sections were washed 2–3 times with 1X-PBS, permeabilized with 1X-PBS containing .1% Triton X-100, blocked with 10% normal goat serum, and incubated with anti-parvalbumin (1:500, Sigma, P3088) overnight at 4° C. Following subsequent washes and incubation with a fluorescent secondary antibody for PV immunostaining and fluorescein-labeled Wisteria floribunda lectin (WFA) (1:1000, Vector Laboratories, FL-1351) for 2 hours at room temperature, sections were washed three times with 1X-PBS, counterstained with DAPI and mounted with Hydromount (National Diagnostics, HS-106) and allowed to dry several days at 4° C prior to confocal imaging.

### PV and PNN cell quantification

Images were acquired using a Leica SP8 laser scanning confocal microscope with an oil immersion, 20X objective with .40 numerical aperture. Laser intensity, gain, and pinhole settings were kept constant for all samples. Images were taken through a z-plane (8.5 μm) within the center of the tissue section, containing 20 stacks (0.4 μm/stack) from the dorsal hippocampus. Quantification of PV numbers with and without an associated PNN cells were counted for each image that were acquired from regions of interest (ROI) using 5–8 mice per group, and 1–3 slides per mouse (total of 13–15 images/group). Due to regional differences in PNN intensity, comparable regions were blindly analyzed for each animal.

### Slice preparation

Following treatment with vehicle (saline) or venlafaxine mice were anaesthetized with deep isoflourane inhalation and then rapidly decapitated. The whole brain was subsequently removed and chilled in cold (0° C) sucrose-based cutting artificial cerebrospinal fluid (sACSF) containing (in mM) 252 sucrose; 3 KCL; 2 CaCl2; 2 MgSO_4_;1.25 NaH2PO4; 26 NaHCO3; 10 dextrose and bubbled by 95% O_2_, 5% CO_2_. Hippocampal slices (480 um thick unless otherwise indicated) were cut in horizontal sections from dorsal to ventral brain with a vibratome (Leica, VT1000S). Horizontal sections were then bisected and right and left sided sections randomly placed to reduce potential confounds due to hemispheric differences. In addition, dorsal and ventral most slices were excluded due to potential PNN density differences along this axis. ACSF contained (in mM) NaCl, 132; KCl, 3; CaCl2, 2; MgSO4, 2; NaH2PO4, 1.25; NaHCO3, 26; dextrose 10; and saturated with 95% O2, 5% CO2 at 26° C. Slices were incubated for at least 120 minutes before being moved to the recording chamber.

### Local field potential (LFP) recordings

Low resistance glass microelectrodes (approximately 150K tip resistance) were used for LFP recordings of gamma frequency oscillations. Electrodes were filled with 1M NaCl in 1% agar, which prevents leakage of the electrode solution that could potentially alter conditions at the recording site. The recordings were done in a submerged chamber, and slices were perfused on both sides at a high flow rate (20 ml /min). Recordings were performed in CA1 stratum oriens proximal to CA2. For exposure of slices to carbachol, infusion was switched from ACSF to ACSF containing carbachol at 40 μM. After 150 seconds, the next 315 seconds were used for analysis (see next section).

### Local Field Potential (LFP) Analysis

The LFP signal was analyzed in 315s blocks, beginning 150s after the onset of carbachol perfusion. 60Hz line noise and harmonics were notch-filtered in pClamp 11 for subsequent analysis with a custom MATLAB algorithm. A Gaussian FIR band-pass filter with corrected phase delay was applied between 1–1000Hz, after which additional band-pass filters were applied for theta (4–12Hz), low gamma (25–55Hz), and high gamma (65–85Hz) ranges. The power was computed by integrating the entire 315s band-pass filtered signals.

### Statistical analysis

Data was entered into a Graph Pad Prism program and statistical analysis performed using Student’s t-test for 2 group comparisons or ANOVA for comparisons of more than 2 groups. Significance was set at p</ = 0.05.

## Results

### I. MMP-9 protein levels are increased, and PNN immunoreactivity reduced, in venlafaxine treated mice

Previous studies have linked increased expression of molecules including brain derived neurotrophic factor (BDNF) to monoamine reuptake inhibitor effects (24). This work does not, however, preclude a significant role for additional molecular effectors of antidepressant efficacy. Previously published data has shown that venlafaxine is linked to increased mRNA expression of MMP-9 in the rat brain (20). With respect to the potential to disrupt PV to pyramidal cell input, which would likely have a profound influence on widespread population activity, MMP-9 has also been shown to diminish PNN integrity (16, 18). Since MMP-9 has been linked to reductions in PNN levels in the setting of both FXS (18) and light reintroduction following dark exposure (16), we examined PNN levels in control and venlafaxine treated mice.

Hippocampal lysates were analyzed by ELISA for levels of MMP-9. As shown in figure 1A, venlafaxine treatment significantly increased hippocampal concentrations of this enzyme (n = 7–8; p = 0.0129; Student’s t-test). For results shown in figure 1B, hippocampal sections were stained for both PV and PNN immunoreactivity and the ratio of PV positive to PNN positive cells quantified. In venlafaxine treated animals, the ratio was significantly increased (n = 8; N = 13–14 *p* = 0.023; Student’s t-test). PV numbers were not increased (control 8.15 +/− 0.87; venlafaxine 7.93 +/− 0.85; p = 0.84), suggesting that PNN positivity was selectively reduced. Hippocampal PV and PNN staining for slice subsections are shown in figures 1C and D. The scale bar represents 75 μM and representative PNN enwrapped PV cells are indicated by arrows.

**I.**
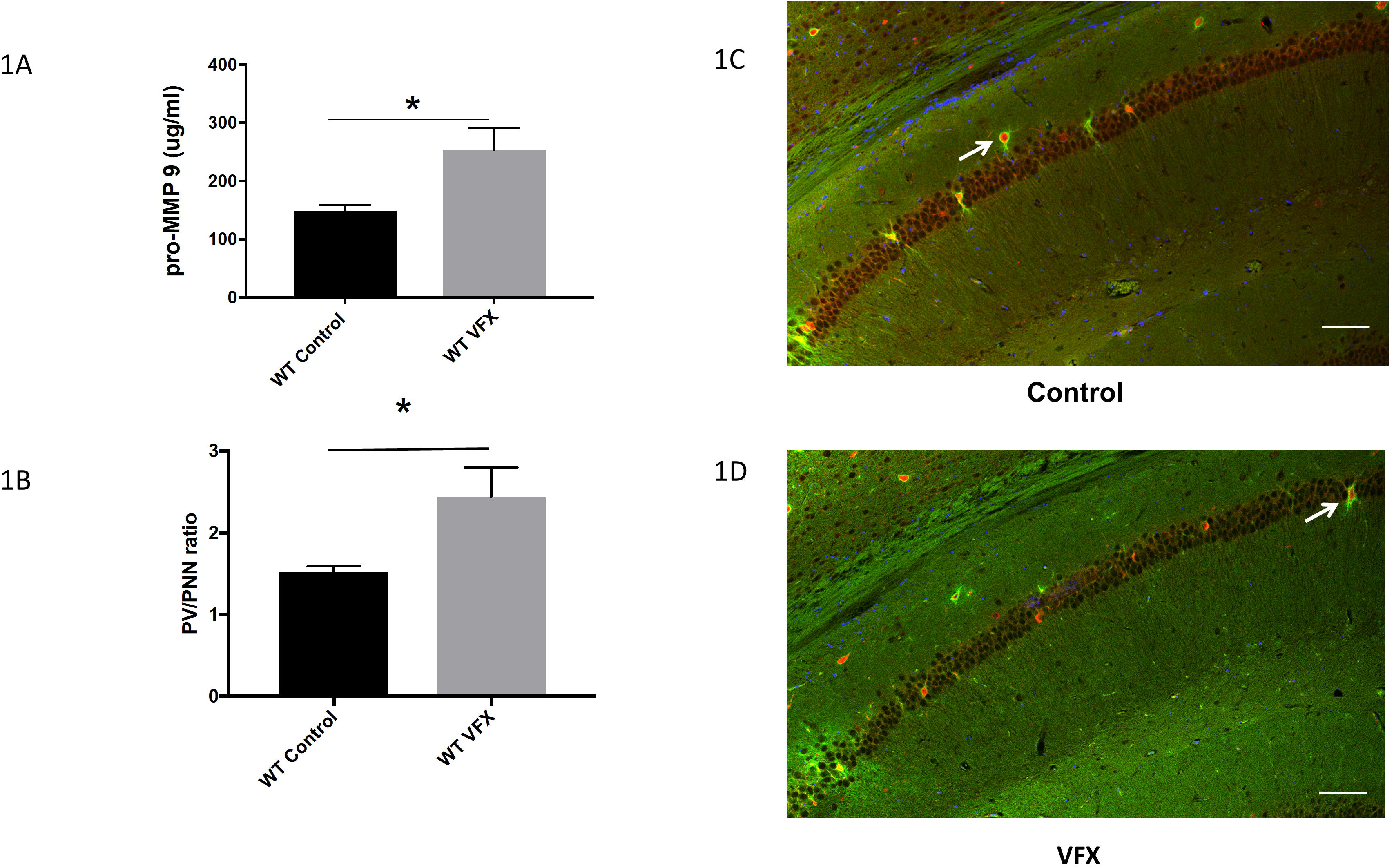
MMP-9 levels and PNN immunoreactivity in hippocampi from venlafaxine treated mice. **1A**) MMP-9 concentrations in brain lysates from 8 control and 7 venlafaxine (VFX) treated mice. Venlafaxine treatment significantly increased hippocampal concentrations of this enzyme (Control 149.4 +/− 9.9; VFX 253.8 +/− 37.3; p = 0.0129; Student’s t-test). **2B**) A quantitative analysis of PV/PNN ratios from control and venlafaxine treated animals (n = 8 animals and N = 13–14 slides for each group) revealed a statistically significant increase in the venlafaxine group (*p* = 0.023; Student’s t-test). PV numbers were not increased (control 8.15 +/− 0.87; venlafaxine 7.93 +/− 0.85; p = 0.84; not shown graphically), suggesting that PNN positivity was selectively reduced. **2C**) PV (red) and PNN (green) immunoreactivity in murine hippocampi in slices from a control and venlafaxine treated animal as indicated. The scale bar represents 75 μM and representative PNN enwrapped PV cells are indicated by arrows.

### II. Venlafaxine increases the power of carbachol-induced gamma activity in ex vivo hippocampal slices

The hippocampus receives cholinergic input from the medial septum-diagonal band of Broca and this input is thought to be an important effector of physiological gamma (25). In isolated hippocampal slices, the cholinergic agonist carbachol is widely used to induce gamma (25). Because chondroitinase mediated PNN degradation has been linked to increased gamma in the visual cortex, and because alterations in gamma may be of relevance to the depressive phenotype (7), we examined the possibility that venlafaxine treatment could impact this rhythm. In figure 2A we show representative local field potential (LFP) recordings between treatments, as well as filtered signals in the theta (4–12Hz) and low and high gamma ranges (25–55 and 65–85Hz respectively). In figure 2B we show there is a statistically significant increase in low and high gamma power for the venlafaxine treated animals (*p* = 0.008 and 0.05 respectively; Student’s t-test; n = 4–6 animals and 1–2 slices per animal for each group). In figure 2C, we show the ratio of post- to pre- carbachol gamma power for each slice. This approach may partially address potential confounds introduced by differences in slice viability or electrode placement. Again we detect a significant difference between slices from wild type saline and venlafaxine treated animals in low gamma (p = 0.0246) and a tendency towards significance in high gamma (p = 0.084).

**II.**
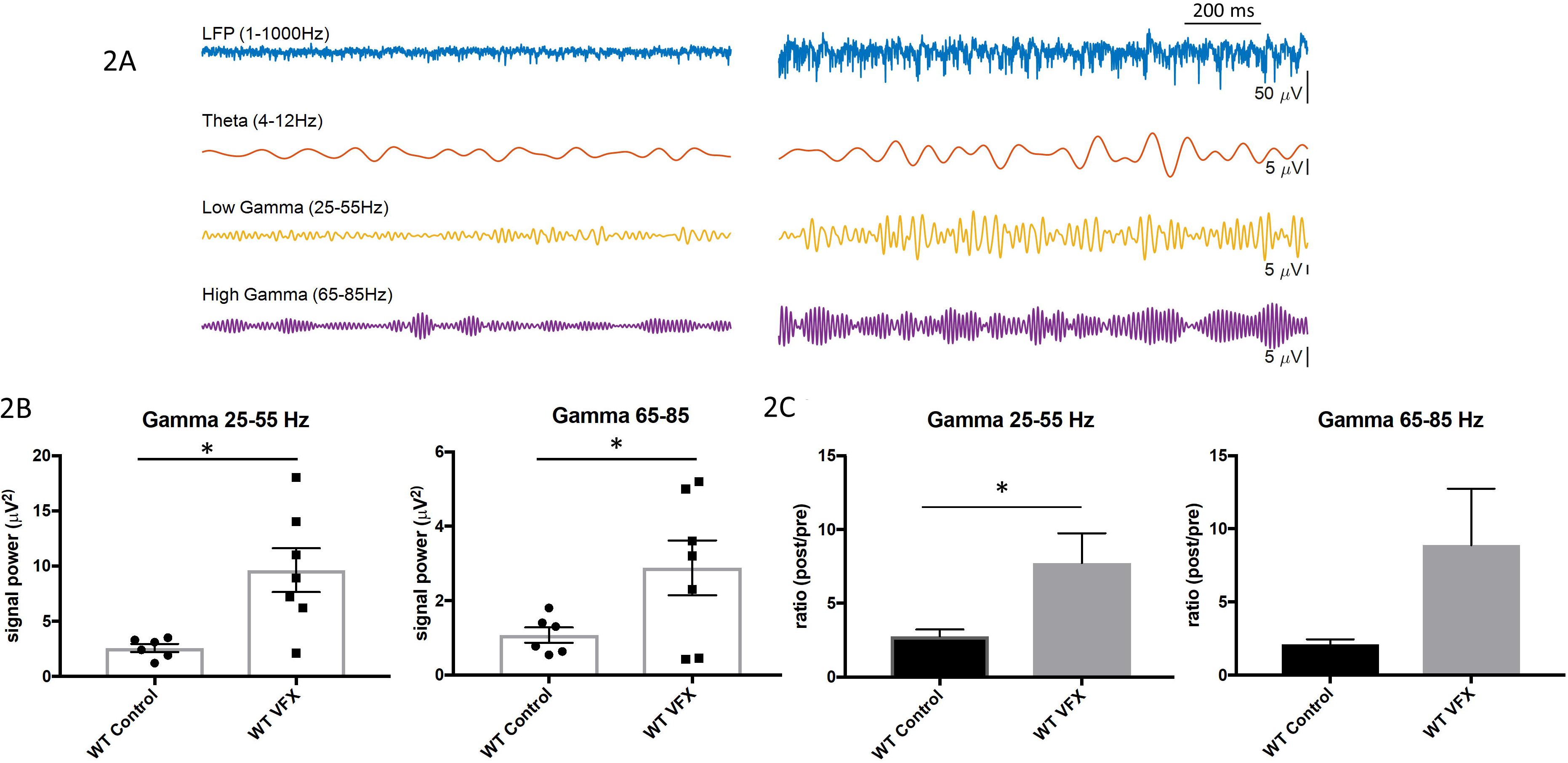
Venlafaxine increases the power of carbachol-induced gamma activity in ex vivo hippocampal slices. **2A**) Representative local field potential (LFP) recordings from slices obtained from a saline or venlafaxine treated animal. The LFP was filtered in the theta (4–8Hz), low and high gamma (25–55Hz and 65–85Hz respectively) ranges. **2B**) The average low and high gamma power of a 315s window beginning 150s after onset of carbachol perfusion was compared between treatment. For n = 4–6 animals, 1–2 slices per animal for each group, there is a statistically significant increase in low and high gamma power for the venlafaxine treated animals *(p* = 0.008 and 0.05 respectively; Student’s t test). **2C**), we show the ratio of post- to pre-carbachol gamma power for each slice. Again we detect a significant difference between slices from wild type saline and venlafaxine treated animals in low gamma (p = 0.0246) and a tendency towards significance in high gamma (p = 0.084).

### III. Venlafaxine stimulated changes in PNN integrity are reduced in MMP-9 null mice

MMP-9 has been shown to reduce PNN integrity in mice that experience light reintroduction after deprivation, and also in a mouse model of FXS (16, 18). MMP-9 may directly target PNN components or activate other proteases that do the same. To examine the possibility that this MMP-9 activity contributes to venlafaxine associated changes in the PNN, we treated MMP-9 null mice with saline or venlafaxine beginning at the same age and with the same dose and duration that had been utilized for wild type animals. Of note, previously published work has examined PNN deposition in wild type and MMP-9 knockout mice (18). Though knockouts showed reduced PV/PNN ratios (consistent with increased PNN) at P21, by P30 baseline PV/PNN levels were comparable between groups (18).

Hippocampal PV and PNN staining for slice subsections are shown in figures 3A and B. PNN enwrapped PV cells are indicated by arrows, while a PV cell without an appreciable PNN is noted by the arrowhead. In 3C we show the PNN/PV ratio for both groups, with data from wild type mice again shown for comparison. There is no significant difference between control and venlafaxine treated MMP-9 null mice (1-way ANOVA; Tukey’s post hoc). In addition, the difference between control wild type and MMP-9 null mice is not significant (1-way ANOVA; Tukey’s post hoc). Interestingly however, there was a significant difference in the PNN/PV ratio in the control and venlafaxine treated MMP-9 null mice when these 2 groups were instead compared with Student’s t-test. The difference was reduced in magnitude as compared to the difference in wild type animals (1.6 fold increase in WT; 1.2 fold increase in MMP-9 null), but the persistence of differences in the MMP-9 null animals suggest that additional PNN degrading proteases may have been upregulated by venlafaxine.

**III.**
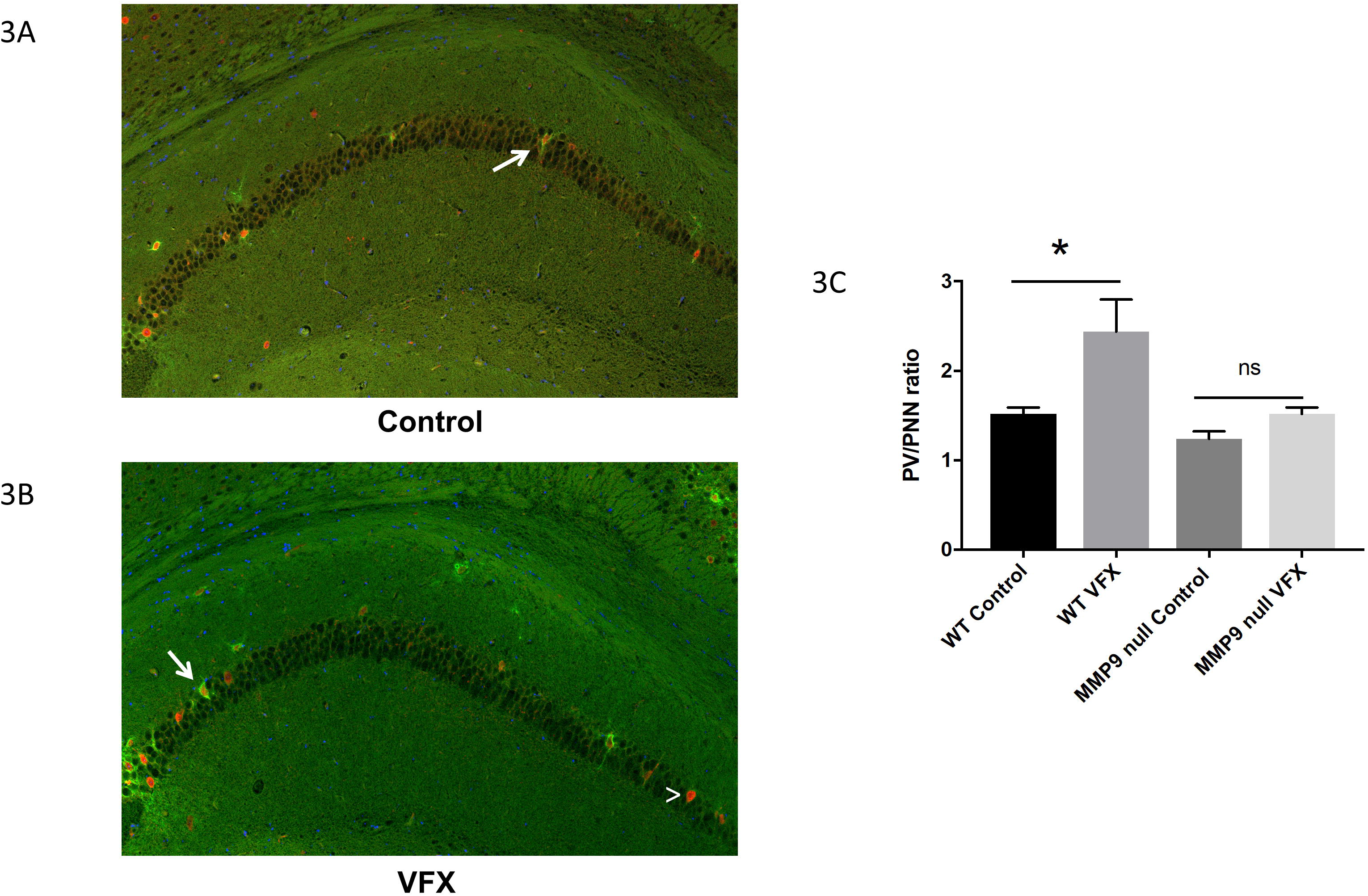
Venlafaxine stimulated changes in PNN integrity are reduced in MMP-9 null mice. **3A-B**) PNN and PV staining in slice subsections from treated and untreated MMP-9 null mice. Representative PNN enwrapped PV cells are indicated by arrows, while a PV cell without an appreciable PNN is noted by the arrowhead. Scale bar represents 75μM. **3C**) PNN/PV ratio for both groups, with data from wild type mice shown for comparison. There is no significant difference between control and venlafaxine treated MMP-9 null mice (1-way ANOVA; Tukey’s *post hoc*). As in wild type animals, the number of PV positive cells was not affected by venlafaxine (control 6.87 +/− 0.79, N = 14; venlafaxine 5.75 +/− 0.59, N = 16; p = 0.26). In addition, the difference between PV/PNN ratios in control wild type and MMP-9 null mice is not significant (1-way ANOVA; Tukey’s *post hoc*).

### IV. Venlafaxine does not significantly increase the power of carbachol-induced gamma activity in ex vivo hippocampal slices from MMP-9 knockout mice

Given that venlafaxine stimulated a lesser fold reduction of PNN integrity in MMP-9 nulls as compared to wild type, we also examined carbachol-stimulated gamma in saline and venlafaxine animals on the MMP-9 null background. Previous studies have examined baseline neurotransmission in these animals and observed that this was not impaired (26). Similarly MMP-9 inhibitors have no effects on baseline properties of neurotransmission and spine size or morphology [reviewed in (27)].

In figure 4A we show representative LFP recordings and filtered signals in the theta, low and high gamma ranges from slices obtained from saline or venlafaxine-treated MMP-9 knockout animals. In 4B we show that in MMP9 knockout mice, we observed no significant increase in low or high gamma power with venlafaxine treatment (Student’s t test). Wild type mouse results (Figure 2B) shown again for comparison. In figure 4C we show post to pre carbachol ratio data in slices from saline and venlafaxine treated mice, with wild type results (Figure 2C) shown again for comparison. The post to pre carbachol ratio in low or high gamma is not significantly different between control and treatment in MMP-9 null animals.

**IV.**
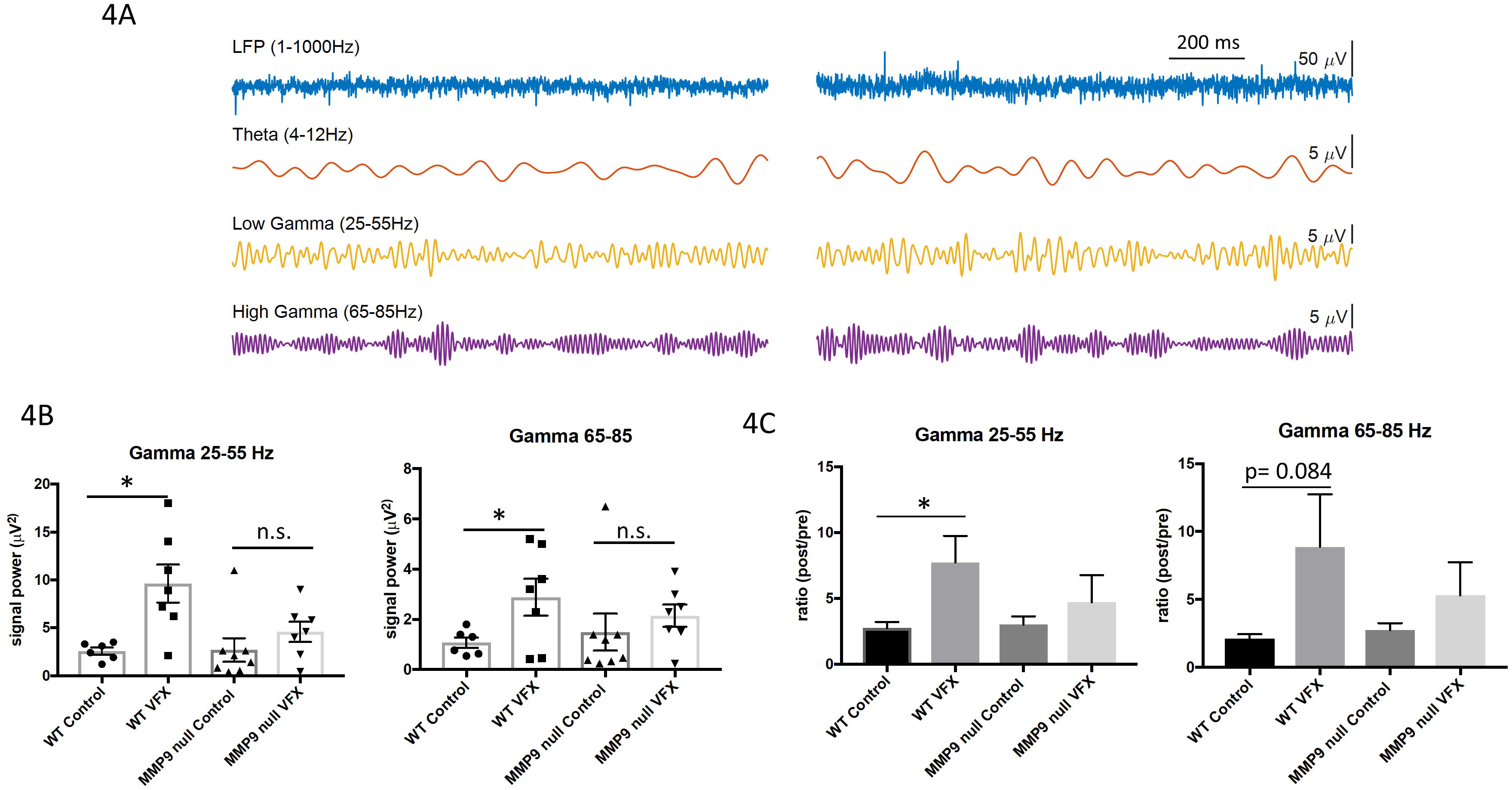
Venlafaxine does not significantly increase the power of carbachol-induced gamma activity in ex vivo hippocampal slices from MMP-9 knockout mice. **4A**) Representative LFP recordings and filtered data from slices obtained from saline or venlafaxine treated MMP-9 null animals. **4B**) Analyses of average low and high gamma power revealed no statistically significant increase due to venlafaxine treatment in MMP-9 null animals (Student’s t test). Wild type mouse results (Figure 2B) shown again for comparison. **4C**) Post- to pre-carbachol ratio in slices from saline and venlafaxine treated mice, with wild type results (Figure 2C) shown again for comparison. The post to pre carbachol ratio in low or high gamma is not significantly different between control and treatment in MMP-9 null animals.

### V. Schematic overview

In figure 5 we show a hypothetical overview by which venlafaxine can influence PNN integrity and gamma power. First, hippocampal levels of serotonin and norepinephrine are increased in the background of venlafaxine treatment. These monoamines interact with G protein coupled receptors expressed on neurons and glia. Gαs protein coupled monoamine receptors in particular, including 5HT7 and the β1 adrenergic receptor (ADRB1), have been linked to increased MMP-9 expression [(19), Alaiyed S. *et al*., unpublished observations]. Increased levels of MMP-9 can in turn target PNN components and/or activate additional proteases that do the same. Reduced PNN integrity, likely localized to regions where monoaminergic transmission typically occurs, can in turn disrupt PV mediated inhibition of pyramidal cell activity with consequent effects on gamma power.

**V.**
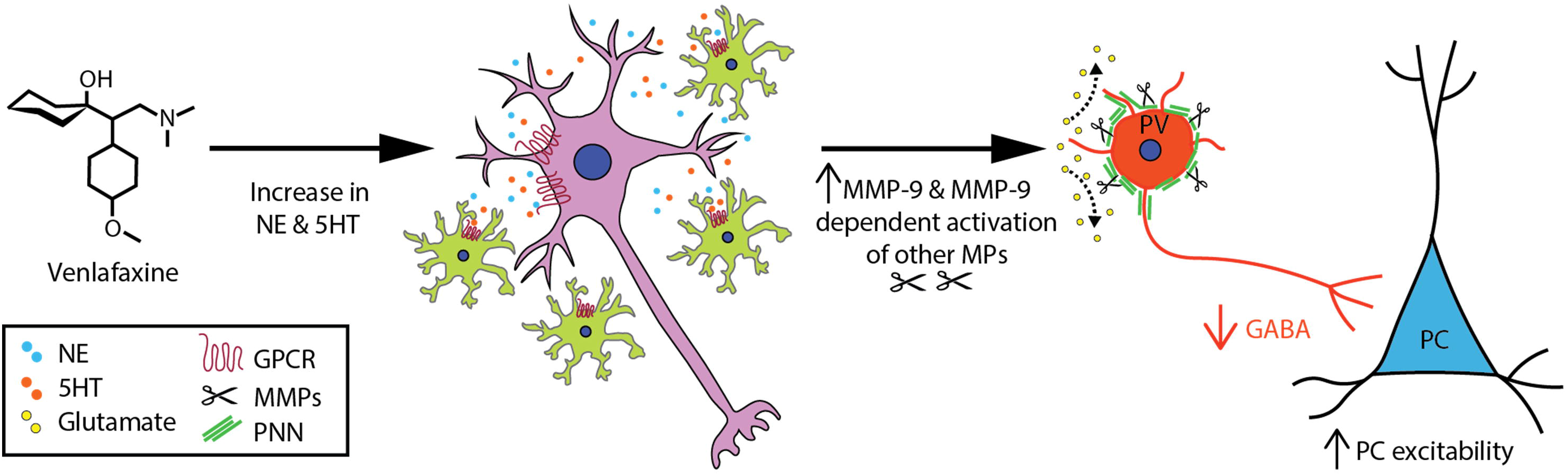
Hypothetical Overview. Hippocampal levels of serotonin and norepinephrine are increased in response to venlafaxine treatment (far left). These monoamines interact with G-protein coupled receptors expressed on neurons (purple) and non-neuronal cells such as microglia (green) to increase expression of MMP-9. Increased levels of MMP-9 can in turn target PNN components and/or activate additional metalloproteases (MPs) that also target PNN substrates. Reduced PNN integrity can in turn disrupt PV-mediated inhibition of pyramidal cell activity with consequent effects on gamma power.

## Discussion

In the present study, we find that chronic treatment of mice with the serotonin/norepinephrine reuptake inhibitor venlafaxine concomitantly increases PNN proteolysis and hippocampal gamma power. We further observe that venlafaxine enhanced gamma power is significantly reduced in mice that are deficient in MMP-9, a family member that is expressed in response to neuronal activity (28) and monoamine exposure (19).

Prior studies have observed regional alterations in gamma power in association with major depressive illness and bipolar disorder (29), as well as increases in gamma with spontaneous remission or selective serotonin reuptake inhibitor (SSRI) treatment in humans and rodent models (6, 30). However, the role of PNN-degrading MMPs has not previously been explored as a mediator of enhanced gamma or antidepressant efficacy. Of interest, the SSRI fluoxetine has been linked to reductions in PNN integrity (13), and conversely, a recent study observed increased hippocampal PNN deposition in a chronic restraint stress rat model of depression (14). Moreover, exogenous chondroitinase, which degrades PNN components, improved stress-associated cognitive deficits in this model (14). In contrast, in mice that are deficient in the PNN protein neurocan, mania-like symptomatology with increased sucrose preference and reduced immobility in the forced swim test are observed (31). Also of relevance are human studies that have examined the PNN in the background of depression and observed alterations including PNN sulfation patterns that render the PNN more resistant to proteolysis (32).

Changes in PNN integrity have the potential to influence oscillatory activity in varied brain regions. This specialized extracellular matrix (ECM) is predominantly localized to PV expressing fast spiking GABA releasing interneurons, which are in turn critical to rhythmic oscillations in the gamma frequency range. At the single cell level, disruption of the PNN has been linked to reduced glutamateric input to PV expressing interneurons (11) through mechanisms that may include increased lateral diffusion of GluAs and presynaptically released glutamate (10). Similarly, genetic deletion of the PNN component brevican has been associated with altered intrinsic properties and reduced synaptic input to PV interneurons through changes in potassium channel localization and levels of GluAs (33). Of interest, antidepressants including venlafaxine have been linked to an increased risk of seizures and disinhibition could contribute to the same.

In terms of effects on population events to which PV activity contributes, PNN disruption has been shown to increase gamma power in the visual cortex (12). Indeed, chondroitinase removal of PNNs was shown to enhance both spontaneous gamma power in the 30–40 Hz range and monocular deprivation-linked gamma activity (12). These effects may seem at odds with the critical role that PV interneurons play in gamma and ripple generation, and with data linking knock down of GluA1 expression on PV cells to reduced gamma power (34). PNN attenuation, however, as opposed to a complete loss of this structure, could enhance gamma power through PC disinhibition in the background of PV cells that retain their ability to respond to the particularly strong rhythmic excitatory post synaptic potentials arising from pyramidal cells during oscillatory activity (35). Consistent with this are results linking a partial block of GluAs on interneurons to gamma generation (36). In addition, ketamine, which preferentially inhibits GluNs at fast spiking interneurons (37), has been associated with increases in gamma power (38). Similarly, intermittent theta burst transcranial magnetic stimulation reduces PV expression and increases EEG gamma power (39). Furthermore, reducing NMDA mediated drive of cortical interneurons increases pyramidal cell firing rate (37) and modeling studies are also consistent with a causative role for PV deficits in terms of oscillatory power (40).

Despite potential similarities between effects of PNN disruption and models of reduced GABA mediated inhibition, we acknowledge that effects of chondroitinase and monoamine uptake inhibitors could be relatively complex. With respect to the latter, given the non-mutually exclusive potential for MMPs to enhance excitatory neurotransmission through relatively direct effects on PCs (41), future studies are warranted to parse out the relative contribution of PNN dependent and independent mechanisms to MMP associated changes in gamma power. One possibility would be to use PNN sulfation mutants that are relatively more resistant to proteolysis (42). It would also be of interest to evaluate gamma in PNN link protein mutants, which display juvenile plasticity and attenuated PNNs (43).

In terms of the potential for increased gamma to contribute to antidepressant efficacy, we acknowledge that future behavioral studies will needed to address this issue. Gamma changes might simply represent a correlate of antidepressant efficacy. Given that increased gamma is observed not only with spontaneous remission but with improvements on the Hamilton Depression scale (30), however, the association is strong. Moreover, in a mouse model, deficits in gamma are inversely correlated with behavioral despair (40). In addition, gamma activity is thought to improve cognitive features that are impaired in major depressive disorder including short-term memory and attention (22). Of additional interest, depressed patients have poor dream recall as compared to controls (44), and both venlafaxine and increased gamma activity have independently been linked to lucid dreaming or enhanced dream recall (45, 46). An unanswered question, however, relates to whether antidepressant associated changes in gamma activity would be wholly beneficial.

In experiments herein, we focused on carbachol-induced gamma in the hippocampus, a region that has been increasing implicated in MDD (reviewed in (22)). For example, depression can be induced by stress in varied rodent models and the hippocampus is sensitive to stress induced atrophy (22). Moreover, hippocampal dependent functions including explicit memory are impaired with depression (22). The hippocampus is also part of a functional network, which includes regions such as prefrontal cortex, that is dysregulated in MDD (22). Of interest, the hippocampus is also the focus of a study examining kainate-induced gamma in *ex vivo* rat slices following acute or chronic administration of fluoxetine or imipramine (47). Differences in species and animal ages, which influence PNN levels (16, 18), may confound direct comparisons with the present study, as may the use of alternate antidepressants and a different gamma-inducing stimulus. Interestingly, however, while gamma power in chronically treated rats was not increased when examined over a broad range, the power trace for fluoxetine at frequencies below 40 Hz was elevated relative to control and imipramine (47).

Herein we also focused on evaluation of gamma in *ex vivo* slices after chronic antidepressant administration. Monoamine reuptake inhibitors typically require several weeks to show a clinical benefit. We posit that changes in PNN integrity could take days to weeks to develop but future studies will be necessary to address the time course by which venlafaxine influences PNN integrity. Importantly, however, we note that acute treatment of *ex vivo* hippocampal slices with monoamines or serotonin reuptake inhibitors can reduce gamma power suggesting that results described in the present study are likely due to changes that follow chronic as opposed to acute antidepressant exposure (47–49).

While our study also demonstrated increased MMP-9 in the CNS of venlafaxine treated animals as well as MMP-9 dependent diminution of the PNN, we acknowledge that other MMPs may contribute to venlafaxine-associated changes in the PNN. This is supported by the observation that the PV/PNN ratio is also increased in the venlafaxine treated knockouts in one analysis (Student’s t-test versus ANOVA). The observation that ratio differences are more robust in the wild type, however, is not unexpected. As compared to other family members, neurons express high levels of MMP-9 when stimulated with glutamate or monoamines (19, 28). MMP-9 is also rapidly transcribed with neuronal activity and LTP (26, 28), and the latter may be enhanced by monoamines (19). It is thus possible that localized increases in monoamine levels can increase neuronal expression of MMP-9 in a relatively selective manner. Light reintroduction and FXS related changes in fragile X mental retardation protein may do the same (50), and MMP-9 has been implicated in the PNN remodeling that occurs in both settings (16), (18). Once upregulated, MMP-9 could also activate additional metalloproteinases that can in turn process PNN components.

In summary, we have shown that chronic treatment with the monoamine reuptake inhibitor venlafaxine increases PNN proteolysis and gamma power in murine hippocampus. We suggest that future studies are warranted to determine whether PNN proteolysis contributes to antidepressant efficacy. While we acknowledge that not all patients respond to monoamine reuptake inhibitors, these medications are effective in a substantial percentage. The identification of new molecular targets could thus lead to the design of drugs or combination therapies. Our results raise the possibility that preferential targeting of GPCRs which best activate signaling molecules linked to increased MMP expression might be pursued.

## Acknowledgements

We would like to acknowledge outstanding veterinary support from the Department of Comparative Medicine and we would also like to apologize to investigators whose excellent work could not be directly cited due to publisher limits.

